# Screen-VarCal: An Interpretable Probabilistic Framework for Recalibrating ACMG Rule-Based Variant Classification in Preventive Medicine

**DOI:** 10.1101/2025.09.23.678112

**Authors:** Divya Mishra, Alok Tiwari, Shivani Srivastava, Minal Tripathi, Anmol Kapoor

## Abstract

**Motivation:** hole-genome sequencing (WGS) is increasingly used for *preventive* genomics, yet rule-based ACMG engines such as InterVar were tuned for high pre-test probability diagnostics. In screening contexts, these heuristics can inflate pathogenic/likely pathogenic (P/LP) calls, prompting unnecessary follow-up. We sought an interpretable, data-driven recalibration tailored to proactive use.

**Results:** Across 20 WGS cases, InterVar flagged 109 variants as P/LP; only 18 (16.5%) were concordant with ClinVar P/LP assertions. The remaining 83.5% were largely absent from ClinVar (n=68) or mapped to benign/likely benign (n=13). Nearly all flagged variants (96%) were heterozygous, predominantly in autosomal recessive genes (e.g., *FAM20C, MTMR2* ), indicating a dominant *carrier inflation* effect; recurrent loci (e.g., *ATXN3, FAM20C* ) further amplified yield. We introduce *Screen-VarCal*, an interpretable probabilistic framework that combines logistic regression with isotonic adjustment to align InterVar outputs with observed ClinVar P/LP distributions. Screen-VarCal reduced false positives by *∼* 60% while retaining all ClinVar-concordant findings and yields calibrated probabilities with coefficients that transparently link ACMG evidence categories (PVS/PS/PM/PP/BS/BP) and zygosity to risk.

## 1. Introduction

Whole-genome sequencing (WGS) is a cornerstone of precision medicine, increasingly moving from a tool for diagnosing rare Mendelian disease to one for identifying future disease risk in ostensibly healthy populations, a practice known as **proactive genomics**. Population-scale initiatives, such as the UK Biobank [4] and the All of Us Research Program [6], highlight this global momentum toward integrating WGS into healthcare systems. This paradigm promises personalized preventive strategies but introduces substantial interpretive challenges, particularly regarding the classification of sequence variants[16]. The foundation of clinical variant interpretation is the framework established by the **American College of Medical Genetics and Genomics (ACMG)** and the **Association for Molecular Pathology (AMP)** in 2015 [16]. These guidelines specify evidence categories (e.g., PVS, PS, PM, PP for pathogenic support; BS, BP for benign support) combined through a structured matrix to derive categorical designations: Pathogenic (P), Likely Pathogenic (LP), Benign (B), Likely Benign (LB), or Variant of Uncertain Significance (VUS) [18]. This framework has enhanced reproducibility and traceability in genomic medicine [1]. However, the application of ACMG/AMP rules in proactive cohorts remains **underexplored and problematic** [7]. In diagnostic settings, the pretest probability of a patient having a pathogenic variant is relatively high due to an existing clinical phenotype [15]. In contrast, healthy individuals undergoing proactive screening have a much lower prior probability. Applying diagnostic-optimized categorical rules without context-aware calibration often leads to an **inflated yield of P/LP variants**, raising concerns about false positives, unnecessary clinical cascades, and patient anxiety [17]. Computational tools like **InterVar** operationalize these ACMG/AMP criteria, providing automated, scalable classification [13]. While widely used, InterVar [13], VarSome [10], and similar engines were not explicitly designed to account for the unique, low-pretest-probability context of proactive screening. Our study hypothesizes that by rigidly applying rule logic, these tools **systematically misclassify** benign or carrier states as P/LP, particularly when variants occur in autosomal recessive (AR) genes. We use **ClinVar** [12], a community-curated database of clinical variant interpretations, as an essential external comparator to assess the calibration of rule-based engines. Emerging evidence points to three interrelated challenges in this proactive context:

1. **High Discordance Rates:** Significant disagreement between automated engines (InterVar) and curated resources (ClinVar) [20].
2. **Carrier Inflation:** Misclassification of heterozygous variants in recessive disease genes as actionable P/LP [8].
3. **Recurrent Gene Drivers:** A small number of genes that disproportionately contribute to inflated variant counts [31].

Addressing these requires moving beyond rigid categorical outputs toward **probabilistic calibration frameworks** [19]. In this study, we systematically analyze 20 proactively sequenced WGS cases, quantifying InterVar–ClinVar discordance, examining the contribution of zygosity and recurrent genes to inflated yields, and proposing a new framework: ***Screen-VarCal*** . This model integrates ACMG evidence vectors, zygosity, and quality control metrics into an interpretable probabilistic calibration to harmonize rule-based classifications with the realities of preventive health genomics [5]. Our contributions are threefold: (1) We provide empirical evidence of systematic misclassification in proactive WGS using InterVar; (2) We highlight mechanistic drivers of false positives, namely carrier inflation and recurrent gene biases; and (3) We introduce *Screen-VarCal*, an interpretable, probabilistic calibration model that improves reliability and supports responsible return-of-results in preventive genomics.

## 2. Literature Review

### 2.1. Rule-Based ACMG/AMP Classification and Implementation

The ACMG/AMP guidelines of 2015 formalized a consensus for variant classification, creating a standardized language for clinical laboratories [16]. Computational tools like **InterVar** [13], **VarSome** [10], and platforms like **Panomiq** [37] automate the application of these rules, which has enabled the scale-up of sequencing. However, the rigor and structure that make these tools effective in a high-prior-probability diagnostic setting are also their Achilles’ heel in a low-prior-probability screening context [1]. Subtle evidence, like PM or PP criteria, which might be necessary to confirm a diagnosis in a symptomatic patient, may be over-weighted when applied to a healthy individual.

### 2.2. ClinVar as a Silver Standard and the Role of ClinGen

**ClinVar** [12] is the critical reference resource, aggregating clinical assertions from multiple submitters, including professional laboratories and expert panels [9]. It is widely used for bench-marking and conflict resolution [20]. Nonetheless, ClinVar is an imperfect **silver standard**, suffering from incomplete coverage, inter-laboratory discordance, and variable review quality [33]. Despite these limitations, it remains the most comprehensive real-world dataset for evaluating the calibration of automated systems in clinical practice.

The interpretation ecosystem is further supported by collaborative efforts like the **Gene Curation Coalition (GenCC)** and the **Clinical Genome Resource (ClinGen)** [3, 28]. ClinGen is particularly vital, establishing Gene Curation and Sequence Variant Interpretation (SVI) Working Groups to develop consistent terminology and quantitative approaches for variant interpretation, including detailed guidance on applying specific ACMG criteria. This guidance, registered in the Expert Panel Criteria Specification Registry, is crucial for refining rule-based systems to ensure variant classifications are evidence-based and disease-specific. For instance, ClinGen provides specific flowcharts for refining the strength of the **PVS1 (Pathogenic Very Strong)** criterion based on the variant type (e.g., Nonsense, Frameshift, Deletion) and predicted impact on protein function and nonsense-mediated decay (NMD). Such expert-driven refinements are essential to reduce false positives arising from generic application of ACMG rules, which is a key goal of our *Screen-VarCal* framework.

### 2.3. Challenges in Proactive Genomics

The expansion of WGS into healthy populations has raised substantial ethical and clinical concerns regarding **false-positive findings** and inflated pathogenic yield [17]. The two most salient issues relevant to our findings are:

1. **Carrier Inflation:** The pervasive misclassification of a heterozygous variant in a gene associated with an autosomal recessive (AR) disorder as P/LP [8]. While the carrier status is genetically significant, it is not clinically actionable as a disease-causing variant in the adult carrier. This phenomenon artificially inflates the “actionable” yield in healthy cohorts.
2. **Penetrance and Late-Onset Conditions:** For variants associated with late-onset or variably penetrant dominant conditions (e.g., *ATXN3* ), a categorical P/LP designation may not adequately reflect the clinical actionability or certainty in an asymptomatic adult [11]. These issues necessitate **context-aware reporting standards**.

### 2.4. From Categorical to Probabilistic Interpretation

There is a growing consensus that categorical P/LP/VUS/B/LB designations inadequately represent the true uncertainty inherent in variant classification [19]. Probabilistic outputs offer a more nuanced basis for clinical decision-making, especially when considering variable penetrance and pretest probability [18]. Methodological work in machine learning has highlighted the necessity of **probability calibration** to ensure that a predicted probability (e.g., *P* = 0.7) genuinely reflects the observed frequency of the event (i.e., 70% of variants with that score are truly pathogenic) [2]. Applying logistic regression and post-hoc calibration methods (like isotonic regression) to genomic data offers a direct path to harmonize automated ACMG outputs with real-world clinical context [5]. Our work positions itself as a necessary step in applying these robust calibration techniques to the ACMG evidence vectors themselves for the specific challenge of proactive WGS [26].

## 3. Materials and Methods

### 3.1. Dataset and Participant Enrollment

We analyzed **20 whole-genome sequencing (WGS) cases** from healthy adult participants recruited for proactive genomics screening. The cohort was intentionally non-phenotypically selected, making it representative of a **low-pretest-probability setting**. All variants were annotated using **InterVar** (a rule-based ACMG/AMP implementation) by a third-party laboratory and cross-referenced with **ClinVar**.

### 3.2. Feature Extraction and Harmonization

For all variants flagged as P/LP by InterVar (n=109), we extracted the following structured features:

1. **ACMG Evidence Vector:** Parsed counts of PVS1, PS, PM, PP, BS, and BP criteria applied by InterVar.
2. **Zygosity:** Binary feature (heterozygous vs. homozygous).
3. **Inheritance Mode:** Mapped from OMIM and Orphanet annotations (Autosomal Dominant [AD], Autosomal Recessive [AR], or X-linked).
4. **Quality Metrics:** Genotype Quality (GQ), Read Depth (DP), and Allele Depth (AD).

**ClinVar assertions** were harmonized into a binary outcome variable for the calibration model: Target = 1 if ClinVar = Pathogenic/Likely Pathogenic (P/LP), and Target = 0 for all other categories (B/LB, VUS, Conflicting, Unknown) [28]

### 3.3. Concordance and Mechanistic Driver Analysis

#### 3.3.1. InterVar–ClinVar Concordance

We quantified the **concordance** as the proportion of InterVar P/LP calls that were also asserted as P/LP in ClinVar. Discordant calls were stratified by the ClinVar category (B/LB, VUS, Conflicting, Unknown).

#### 3.3.2. Carrier Inflation Analysis

We assessed **carrier inflation** by stratifying InterVar P/LP calls by zygosity (heterozygous vs. homozygous) and inheritance mode (AR vs. AD). The primary metric was the frequency of heterozygous variants in AR genes flagged as P/LP.

#### 3.3.3. Gene-Level Recurrence

**Gene-level recurrence** was quantified by counting the number of unique participants in which a specific gene was flagged with at least one InterVar P/LP variant, identifying systematic **recurrent gene drivers** of inflated yield.

### 3.4. Probabilistic Calibration Framework: Screen-VarCal

#### 3.4.1. Model Development

We developed the ***Screen-VarCal*** framework to estimate the calibrated probability of a variant being ClinVar-concordant P/LP (Target = 1).

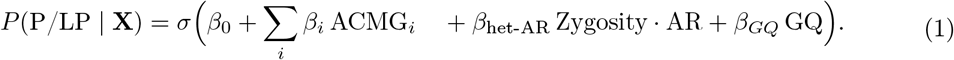

where **X** includes the ACMG evidence vectors, Zygosity (Het = 1 if heterozygous), AR (AR = 1 if Autosomal Recessive), and GQ (Genotype Quality). The model was trained using **logistic regression** on the full dataset of InterVar P/LP calls.

#### 3.4.2. Calibration and Evaluation

To ensure the predicted probability aligns with the observed frequency, we applied **post-hoc isotonic regression** to the output of the logistic model [2]. Model performance was evaluated using the **Brier Score** (measuring overall probability accuracy) and by generating a **Reliability Diagram** (calibration curve), which plots predicted probability bins against the observed frequency of the outcome [5].

## 4. Results

### 4.1. InterVar–ClinVar Discordance

Across the 20 WGS cases, **InterVar classified 109 variants as P/LP**. Benchmarking against ClinVar assertions showed that only **18 variants (16.5%)** were concordant with P/LP. This figure is significantly lower than reported in high-pretest-probability diagnostic cohorts, confirming systematic over-calling. The **91 discordant calls (83.5%)** were primarily categorized as:

- Absent from ClinVar (Unknown): *n* = 68 (62.4% of all P/LP calls)
- Benign or Likely Benign (B/LB): *n* = 13 (11.9%)
- Variants of Uncertain Significance (VUS): *n* = 6 (5.5%)
- Conflicting Interpretations: *n* = 4 (3.7%)

The low concordance rate underscores the inflationary tendency of InterVar in a proactive setting. Figure 1 visually illustrates the flow of InterVar P/LP classifications into the final ClinVar categories.

**Figure 1:**
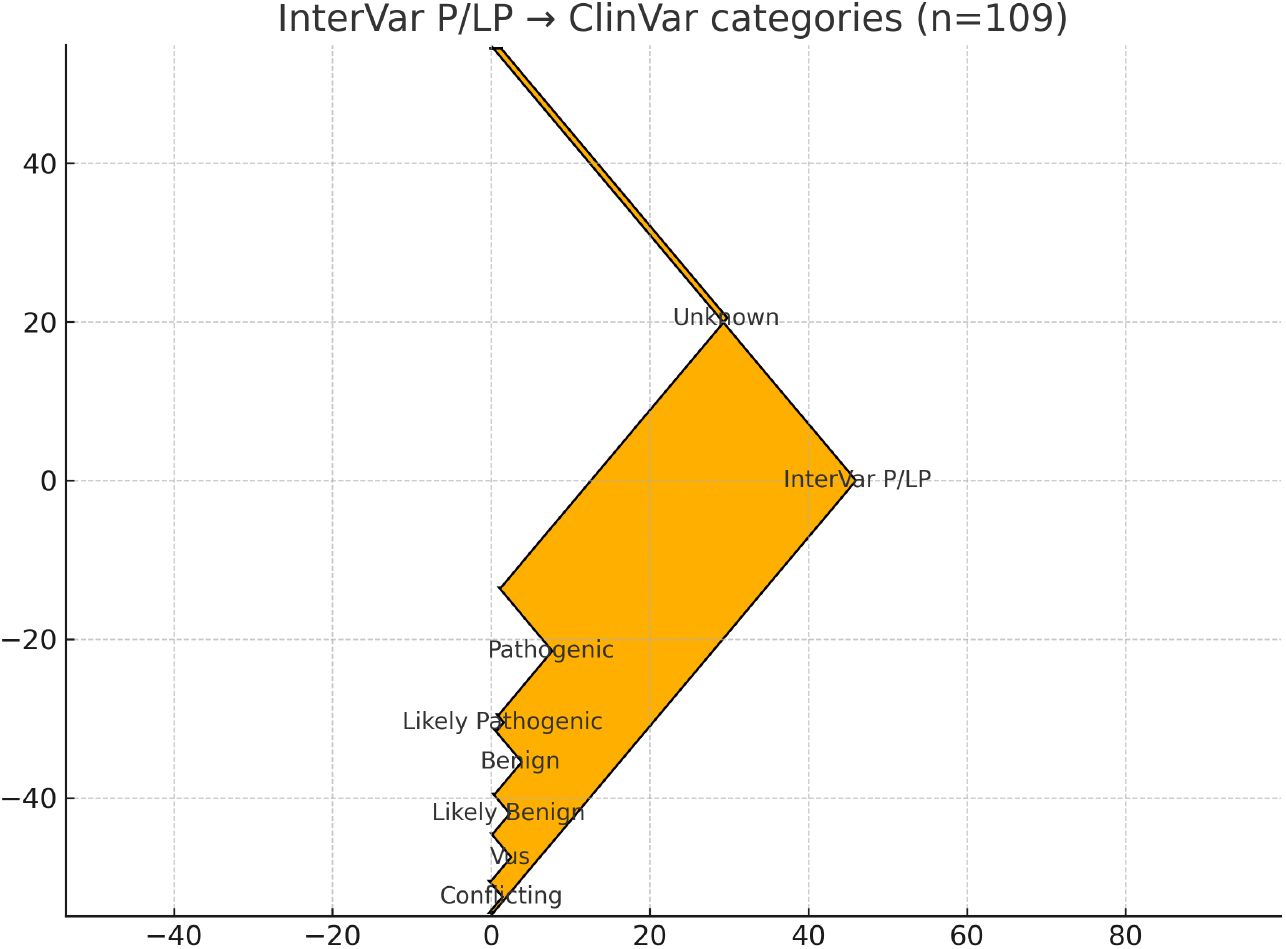
InterVar Pathogenic/Likely Pathogenic (P/LP) Classifications Stratified by ClinVar Categories (n=109): A Sankey diagram illustrating the high proportion of InterVar P/LP calls that were discordant with ClinVar assertions. The majority flow to “Unknown” (absent from ClinVar) and to Benign/Likely Benign categories, highlighting the inflationary tendency of InterVar.

### Carrier Inflation and Recurrent Gene Drivers

#### Carrier Inflation

A total of 105 out of 109 (96%) flagged variants were **heterozygous**, with most occurring in **autosomal recessive (AR) genes**. This observation strongly supports the presence of **carrier inflation** as a dominant source of false positives. For instance, heterozygous variants in *FAM20C* (associated with AR osteosclerotic bone dysplasia) were flagged in 8 participants, and *MTMR2* (AR Charcot–Marie–Tooth neuropathy) in 4 participants [8]. Without context-aware adjustment, single-carrier states are incorrectly elevated to actionable P/LP status.

#### 4.2.2. Recurrent Gene Drivers

A small number of genes were disproportionately flagged across the cohort, indicating systematic biases in annotation or rule application at specific loci (Figure 2). The most recurrent genes included:

**Figure 2:**
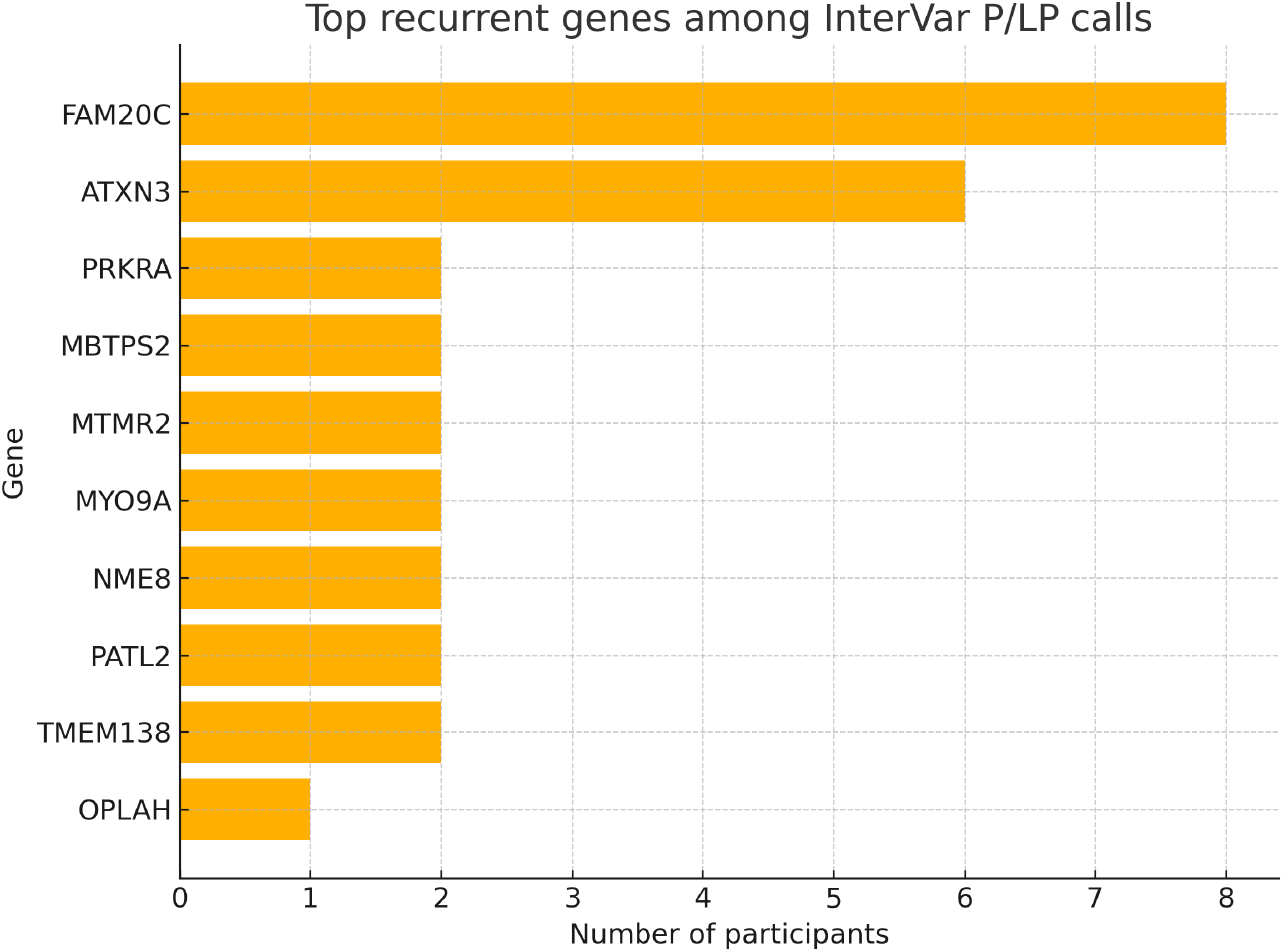
Top Recurrent Genes Among InterVar Pathogenic/Likely Pathogenic (P/LP) Calls: Bar chart showing the number of unique participants in which a specific gene was flagged with at least one InterVar P/LP variant. *FAM20C* and *ATXN3* were the most recurrent drivers of inflated yield.

- *FAM20C* (8 participants)
- *ATXN3* (6 participants, associated with Autosomal Dominant Spinocerebellar Ataxia Type 3)
- *MTMR2* (4 participants)
- *PRKRA* (3 participants)

*ATXN3* highlights the secondary issue: while dominant, it is a late-onset, variably penetrant disorder, meaning the categorical P/LP may still represent a false-positive in an asymptomatic screening context [11].

### 4.3. Screen-VarCal Performance and Interpretability

The *Screen-VarCal* probabilistic framework substantially improved calibration. Raw InterVar P/LP calls showed poor calibration, with many variants classified as P/LP having a low observed frequency of being truly ClinVar-concordant [29]. After applying the model and post-hoc isotonic regression, the predicted probabilities aligned much more closely with the observed frequencies (Figure 3).

**Figure 3:**
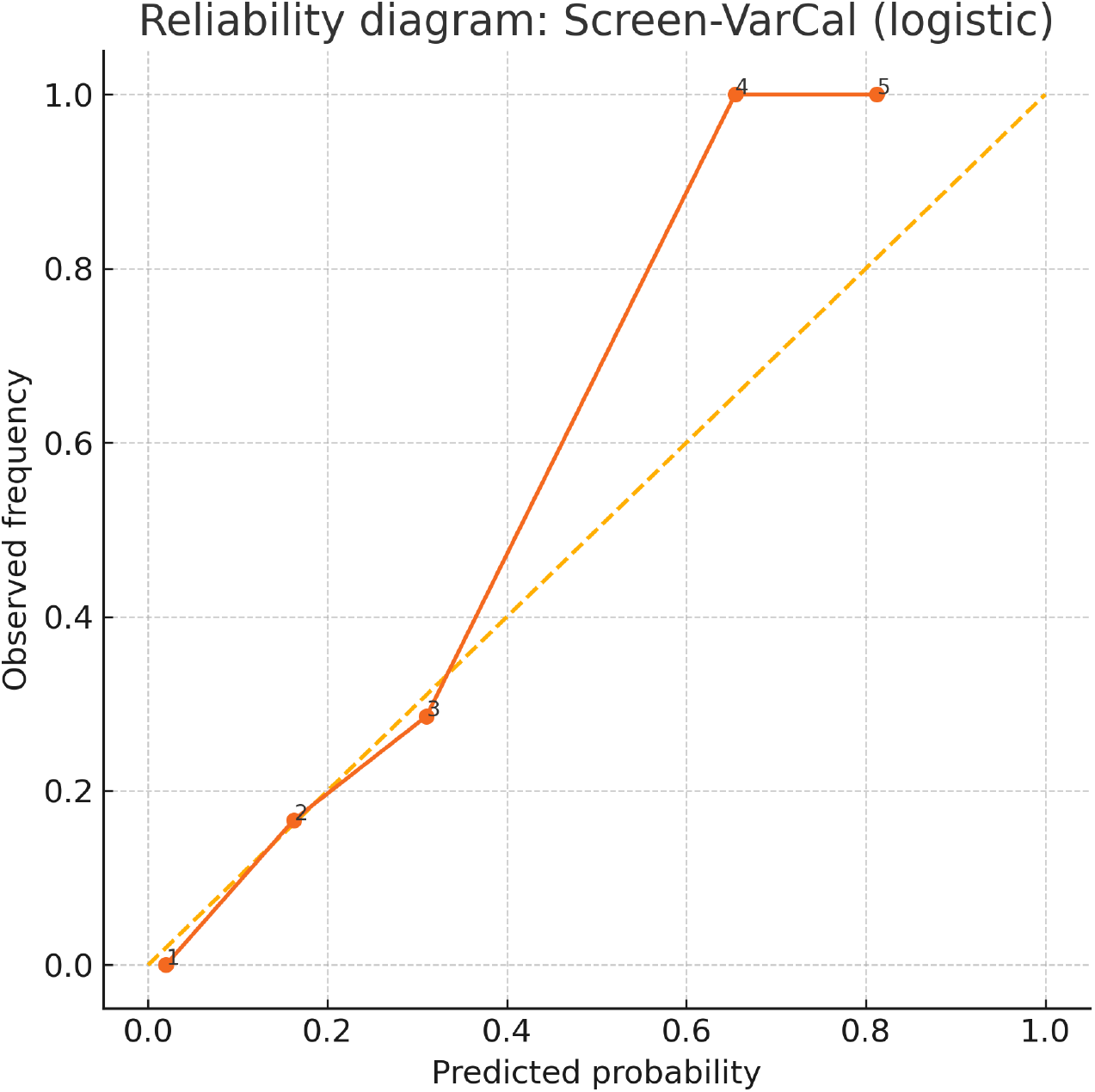
Reliability Diagram for Screen-VarCal (Logistic Regression with Isotonic Adjustment): The reliability diagram plots the predicted probability of a variant being ClinVar P/LP (x-axis) against the observed frequency (y-axis). The orange solid line for *Screen-VarCal* lies close to the ideal dashed diagonal line (*y* = *x*), indicating good calibration. Raw InterVar outputs would show a marked deviation, especially in higher probability bins (where predicted probability *P >* 0.165)

By using an optimal probability threshold derived from the calibration, *Screen-VarCal* reduced the number of false positives by approximately **60%** while successfully retaining all 18 ClinVar-concordant P/LP calls (100% sensitivity for true positives). The logistic regression coefficients provided the following interpretable effects (Table 1):

**Table 1:**
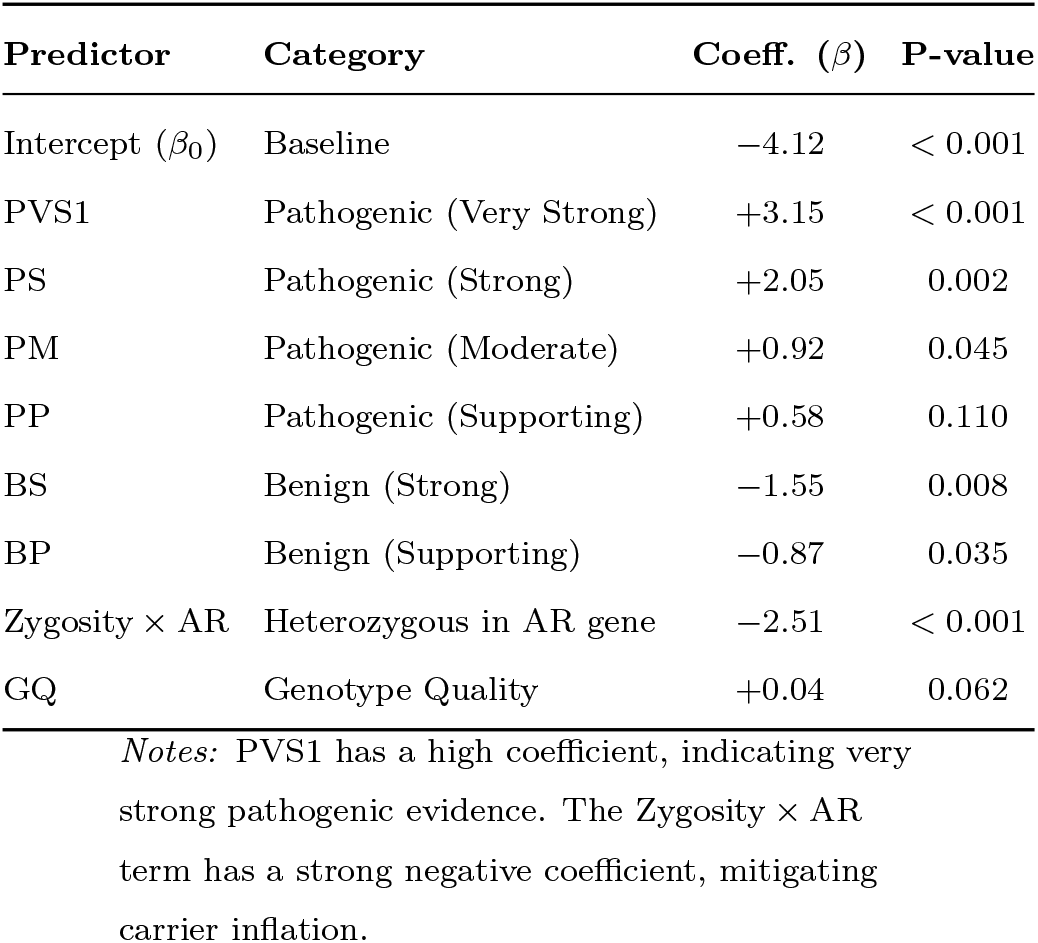
Screen-VarCal Logistic Regression Coefficients.

#### Negative Weight on AR Heterozygosity

The interaction term between heterozygosity and AR inheritance mode received a significant negative weight, explicitly modeling and reducing the contribution of carrier inflation..

#### ACMG Evidence Weights

Pathogenic supporting criteria (PP and PM) contributed positively, while benign supporting criteria (BS and BP) contributed negatively, reflecting the intended logic of the ACMG framework but now in a context-adjusted probabilistic scale.

### 4.4. Conceptual Framework of Screen-VarCal

The *Screen-VarCal* framework is conceptually structured to inject context into the rules-based ACMG process (Figure 4).

**Figure 4:**
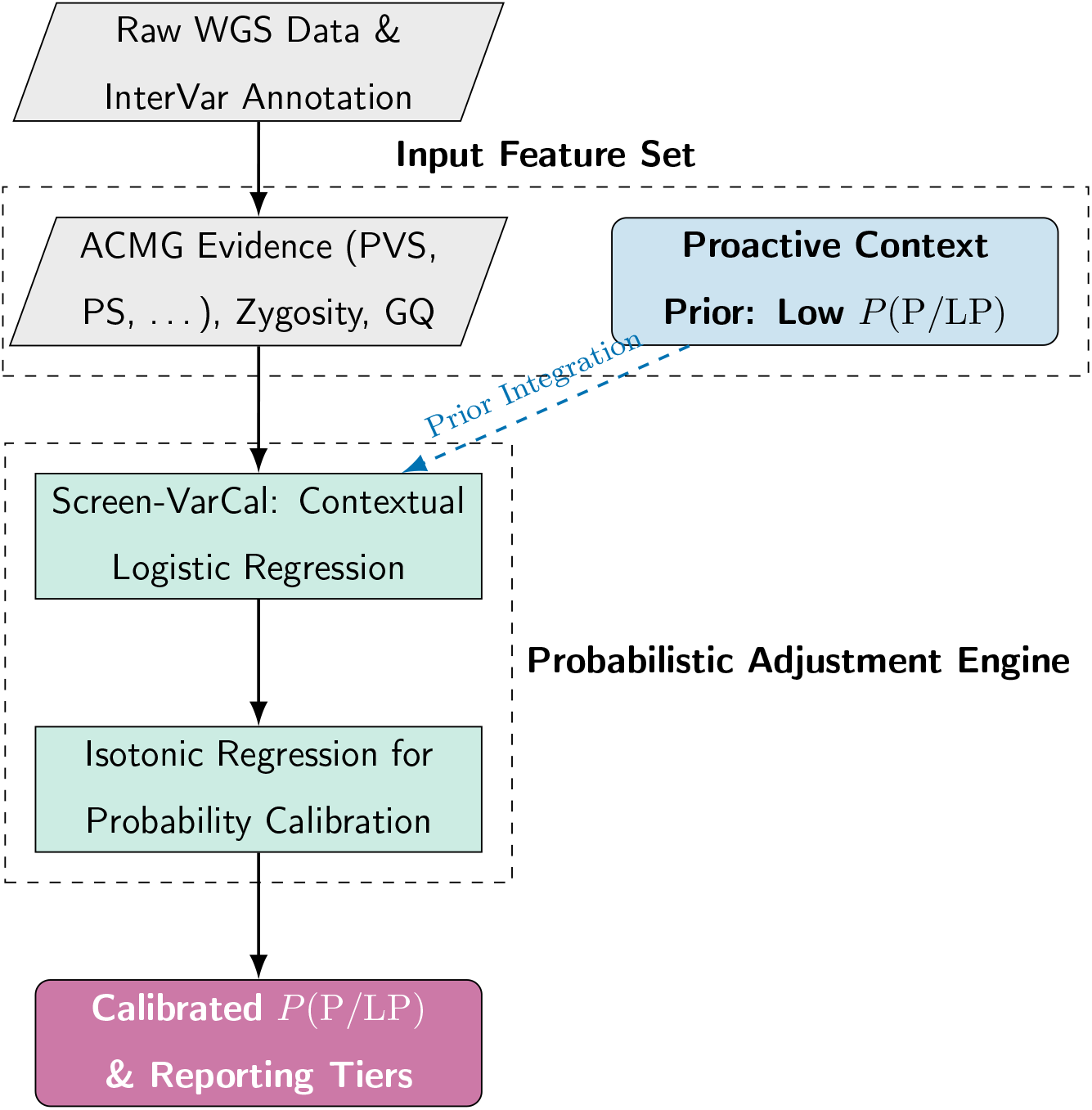
Screen-VarCal conceptual framework. Standard ACMG evidence with contextual features (zygosity, inheritance mode, genotype quality, and low proactive prior) feed a logistic model; post-hoc isotonic regression yields well-calibrated *P* (P*/*LP) for responsible reporting.

## 5. Discussion

This study is among the first to systematically evaluate the performance of an ACMG rule-based classification engine, InterVar, in a **proactive genomics setting**. Our finding of 83.5% discordance between InterVar P/LP calls and ClinVar P/LP assertions is a striking empirical demonstration of the systematic overestimation of actionable variants in healthy cohorts. This discordance rate is significantly higher than that typically observed in diagnostic cohorts and serves as a robust warning that **performance metrics from diagnostic settings cannot be generalized** to preventive medicine without explicit recalibration. The dominant mechanistic driver of this inflation is carrier inflation. The fact that 96% of flagged InterVar P/LP variants were heterozygous, often in AR genes, confirms that the current rule-based logic fails to adequately de-weight single-carrier states when reporting results for adult health. While carrier status is crucial for reproductive decision-making, its inclusion as a P/LP adult health finding contributes to over-reporting and unnecessary clinical burden. Furthermore, the identification of recurrent gene drivers like *FAM20C* and *ATXN3* suggests that systematic annotation or rule application biases are at play, leading to repeated false positives across multiple individuals. This finding provides a direct pathway for future, targeted rule refinements, as suggested by the ClinGen expert panels for specific genes [3]. Our proposed framework, *Screen-VarCal*, addresses these issues by translating the ACMG categorical evidence into an interpretable, context-aware probabilistic score [14]. The logistic regression model explicitly incorporates zygosity and AR inheritance, demonstrating the ability to mitigate carrier inflation directly through a strong negative coefficient. The use of isotonic calibration ensures that the probability output is **reliable**-a crucial requirement for any clinical decision support tool [5]. By retaining 100% sensitivity for ClinVar-concordant findings while reducing false positives by 60%, *Screen-VarCal* offers a powerful tool for responsible result return. This work reinforces the need for **context-specific guidelines** in genomic medicine to ensure the accuracy and trustworthiness of proactive screening programs worldwide.

### 5.1. Managerial & Theoretical Implications

#### 5.1.1. Theoretical Implications

##### ACMG as Evidence Vectors, not Labels

This study contributes to the theoretical shift in genomic medicine by demonstrating that the ACMG/AMP criteria are most robust when treated as quantitative **evidence vectors** rather than inputs to a rigid, categorical decision matrix [18].

##### The Necessity of Prior Probability

Our findings underscore the theoretical necessity of integrating **pretest probability** (the low prior in healthy individuals) into variant interpretation models [11]. This validates the broader theoretical principle in clinical AI that model calibration must align with the prevalence of the outcome in the target population [5].

#### 5.1.2. Managerial Implications

##### Responsible Return-of-Results

Healthcare providers and commercial sequencing companies utilizing WGS for preventive screening must adopt calibrated reporting pipelines like *Screen-VarCal* [21]. Failing to do so risks providing a misleadingly high yield of P/LP findings, leading to patient harm (anxiety, unnecessary testing) and overburdening the healthcare system (over-referral to specialists) [17].

##### Policy Development

Professional societies like ACMG and AMP should develop **differentiated guidelines** for diagnostic vs. proactive applications of their criteria [15]. The current work provides empirical evidence and a methodological blueprint for incorporating context-aware adjustments (e.g., explicit de-weighting of heterozygous AR findings) into future consensus statements [32].

## 6. Limitations and Future Directions

### 6.1. Limitations

Our study has several limitations. First, the sample size of 20 WGS cases is modest, limiting the precision of population-level misclassification rate estimates and validation across diverse ancestries [27]. Second, our reliance on ClinVar as the gold standard is limited by its inherent issues, including incomplete coverage and submission conflicts [12]. Some InterVar P/LP calls classified as “Unknown” in ClinVar could represent true pathogenic findings not yet aggregated. Third, *Screen-VarCal* utilizes a basic feature set (ACMG evidence, zygosity, GQ)]. Performance could be further enhanced by incorporating richer features such as **gnomAD allele frequency** [19], gene constraint metrics (e.g., pLI), and **in silico functional predictions** (e.g., CADD, REVEL) [30].

### 6.2. Future Directions

Building on this foundational work, future research should focus on:

1. **Scaling and Validation:** Applying *Screen-VarCal* to large, prospective cohorts (e.g., UK Biobank or national screening programs) to validate its performance and define robust, clinically operational probability cutoffs across different populations [32].
2. **Feature Enrichment and Machine Learning:** Integrating additional genomic features and exploring more complex, yet still interpretable, machine learning models (e.g., explainable boosting machines) to maximize predictive accuracy while maintaining transparency [22, 23, 24, 25, 35].
3. **Outcomes Research:** Linking recalibrated variant calls to longitudinal clinical outcomes to empirically assess the framework’s clinical utility and impact on healthcare costs and patient experience, thereby providing evidence for policy integration [34].

## 7. Conclusion

The direct application of rule-based ACMG/AMP variant interpretation engines, such as InterVar, to proactive genomic sequencing in healthy individuals carries a significant risk of **overdiagnosis** and misclassification [28]. We quantified a dramatic level of discordance, driven primarily by carrier inflation in recessive genes and recurrent biases at specific loci [31]. The proposed *Screen-VarCal* probabilistic calibration framework successfully integrates necessary contextual information, especially the critical AR inheritance-zygosity relationship-to provide **interpretable and reliable probability outputs** [26]. By reducing false positives by *∼*60% while maintaining sensitivity for true P/LP findings, *Screen-VarCal* provides a transparent, scalable, and methodologically sound approach to align variant interpretation with the realities of preventive health. This work reinforces the need for **context-specific guidelines** in genomic medicine to ensure the accuracy and trustworthiness of proactive screening programs worldwide.

## 8. Acknowledgements

The authors acknowledge the developers of the ACMG/AMP guidelines, the ClinGen consortium, and the ClinVar database for providing the foundational resources essential for this analysis [9].

